# A network-based methodology to reconstruct biodiversity based on interactions with indicator species

**DOI:** 10.1101/2023.10.27.564487

**Authors:** Ilhem Bouderbala, Daniel Fortin, Junior A. Tremblay, Antoine Allard, Patrick Desrosiers

## Abstract

The relationship between species presence, biodiversity reconstruction, and latitudinal gradients is a complex and multifaceted topic that has been the subject of extensive research in ecology. Recent studies have provided valuable insights into the patterns and drivers of these phenomena. Also, with the ongoing decline in biodiversity, there is a need for efficient field monitoring techniques. Indicator species (IS) emerged as a promising tool to monitor diversity because their presence indicates a maximum number of conditionally co-occurring species. We aim to assess the effectiveness of IS for biodiversity reconstruction implicitly based on their co-occurrence with other species through a network-based methodology. The IS are identified based on various network metrics and the likelihood of species’ occurrences is computed based on (1) their conditional occurrence probability with IS and (2) the occurrence probability of IS. We test the approach with field observations of birds in the Côte-Nord region of Québec. From our methodology, the climate latitudinal gradient plays a significant role on the alternation in composition of IS with an almost complete turnover between northern and southern networks. The latitudinal gradient impacts also the nature of the inter-specific interactions with more avoidance relationship toward the Tropics and more cooperation liaisons toward the north. Regarding the effectiveness in the reconstruction of assemblages occurrence, we observe a strong negative correlation (*r ≤ −*0.95) between the percentage of sites occupied and the dissimilarity between the original and the estimated occurrences. More precisely, species must be present in more than 29% and 33% of northern and southern sites to recover well from its co-occurrence with IS. Therefore, it is more challenging to reconstruct biodiversity in communities closet to Tropics due to higher complex interactions and interspecific competition in these areas, which make it more difficult to infer community composition. In conclusion, our method demonstrates that it is possible to predict local species assemblages based on their implicit interactions with local IS. Nevertheless, the relatively low success of less present species illustrates the need for further theoretical development to reconstruct biodiversity, mainly to recover the occurrence of rare species.

## 1 Introduction

Climate change is a significant threat to biodiversity and can potentially disrupt ecosystems’ functioning (Thuiller et al., 2011; Pachauri et al., 2014; Zhang and Liang, 2014). Rising global temperatures and changing weather patterns are causing shifts in habitats and altering the availability of resources necessary for species survival (Duveneck et al., 2014). In particular, in Canada, the rise in the temperature led to the northern migration of thermophilous hard-wood trees, which was replacing boreal conifers (Boulanger and Puigdevall, 2021). In addition to climate change, ecosystems are also impacted by other anthropogenic changes such as land use, pollution, invasive species and the occurrence of extreme climatic events (Masson-Delmotte et al., 2018; Rechkemmer and von Falkenhayn, 2009; Labadie et al., 2021). While climate change plays a significant role in biodiversity loss, land-use changes may be a more vital driver of biodiversity loss in specific ecosystems, particularly in the short term. The cumulative effect of habitat change, occurring through anthropogenic disturbances, with the influence of climate and meteorological conditions, impacts heavily ecosystem functioning (Boulanger et al., 2017; Micheletti et al., 2021). Indeed, the coupled effect of climate change and land use leads to biodiversity alternation mainly caused by species turnover (Pachauri et al., 2014; Cadieux et al., 2020; Bouderbala et al., 2023b). These changes in biodiversity could also exert pressure on agroecosystems and result in a loss of ecosystem services (Kelly and Goulden, 2008; Hillebrand et al., 2010).

The significance of monitoring biodiversity and the development of reconstruction tools has become increasingly crucial (Azeria et al., 2009; Dommain et al., 2020). Monitoring biodiversity allows us to understand the causes and current state of biodiversity loss to identify the main drivers of biodiversity decline or increase (Bouderbala et al., 2023a), such as habitat destruction, pollution, climate change, and invasive species. Moreover, monitoring biodiversity helps plan mitigation strategies and implement effective conservation actions (Stralberg et al., 2015). Furthermore, monitoring biodiversity is necessary to evaluate the effectiveness of conservation measures and adjust them accordingly (Robichaud and Knopff, 2015; Labadie et al., 2023). However, identifying species of many taxa to reconstruct biodiversity needs expertise and could be costly (Kim and Byrne, 2006). Therefore, methods based on IS to represent species assemblages are well used to monitor wildlife conservation and ecological restoration (Dufrêne and Legendre, 1997; Chu et al., 2021). Usually, few IS could be informative to monitor environmental changes and provide signals for plausible ecological shift (Siddig et al., 2016; Chu et al., 2021). To use a fixed set of IS over a large spatiotemporal scale, those species must effectively predict some indicator, such as species richness, despite environmental changes. However, the ongoing changes in climate conditions permanently shape biodiversity patterns and community composition (Santillán et al., 2018; Bouderbala et al., 2023b). Therefore, the predictive potential of IS over a large spatial extent must be studied cautiously.

The statistical methods used to determine indicator species (IS) are evolving rapidly and have been utilized as tools for informing management decisions (Fleming et al., 2020; Chu et al., 2021). A method based on null models of species co-occurrence classification, which identifies sets of species with similar and distinct response patterns to their environment, was developed by Azeria et al. (2009) and Terrigeol et al. (2022). Fleming et al. (2020) introduced conceptual models of IS that include direct relationships between an IS, ecosystem change drivers, and latent processes and variables. Additionally, Chu et al. (2021) developed an approach for incorporating species-conditional co-occurrence into models used to select marine IS. The targets of null models is to describe the extent between random patterns and structured co-occurrence processes (Veech, 2013). However, the metrics and algorithms used for randomizing species presence-absence data might differ, leading to different statistical and computational properties (Rivest and Ebouele, 2020), such as varying Type I and II error rates (Gotelli, 2000). A Type I error occurs when a randomly associated pair of species is incorrectly identified as being either positively or negatively associated, while a Type II error occurs when a truly positively/negatively associated pair is incorrectly identified as being randomly associated (Veech, 2013). One source of Type I and Type II errors is when an algorithm does not randomize the species presence-absence matrix in an unbiased way. In this article, we use a probabilistic species cooccurrence model that does not rely on any data randomization, resulting in a very low Type I error rate and maintaining high power with a low Type II error rate (Veech, 2013).

In this work, we proposed a network-based methodology to assess the reconstruction of biodiversity from interactions with indicator species (IS) along the latitudinal gradient. We quantified the effectiveness of this reconstruction by comparing the estimated assemblage occurrence, using our approach, to the original occurrence based on: (1) the co-occurrence with the local IS of the assemblage and (2) modeling the occurrence of the IS. Furthermore, we emphasized the conditions under which biodiversity could be reconstructed from interactions with local IS.. Our analysis focused on bird species assemblages in the Côte-Nord region of Québec, Canada, covering an area of 114,118 km2. We developed a model to identify IS based on a probabilistic species co-occurrence analysis (Veech, 2013) along a latitudinal gradient (Lurgi et al., 2020). The conception of latitudinal co-occurrence networks offers a unique opportunity to identify changes in IS composition along the latitudinal gradient. The identified latitudinal climate clusters represented spatial variations in short-term climate conditions. We constructed latitudinal co-occurrence networks for each climate cluster and identified the indicator species within each network based on network metrics evaluated through co-occurrence significance. The latitudinal analysis provided insights into how biotic interactions are changing, helping us understand how shifts in climate conditions will alter community composition (Graham and Grimm, 1990) and change the strength of interspecific interactions (Parmesan and Yohe, 2003; Harley, 2011). Our primary goal is to estimate the probability of occurrence for all non-indicator species in the assemblage, conditional on the environmental conditions. The model implicitly includes the structure of dependence on the IS by incorporating the probability of presence, conditional on the presence or absence of the associated IS

## 2 Network-based community occurrence model

### 2.1 Framework

Through our framework, we first modelled the occurrence of the selected IS based on a single-species model as a function of the climate variables and the occurrence of the other IS in the same climate region. The second step consisted in calculating the occurrence probability of species in the pool conditional to the presence and the absence of the IS. Finally, we combined the two precedent probabilities to estimate the community occurrence. We divided our approach into six steps (see Fig. 1).

**Figure 1.**
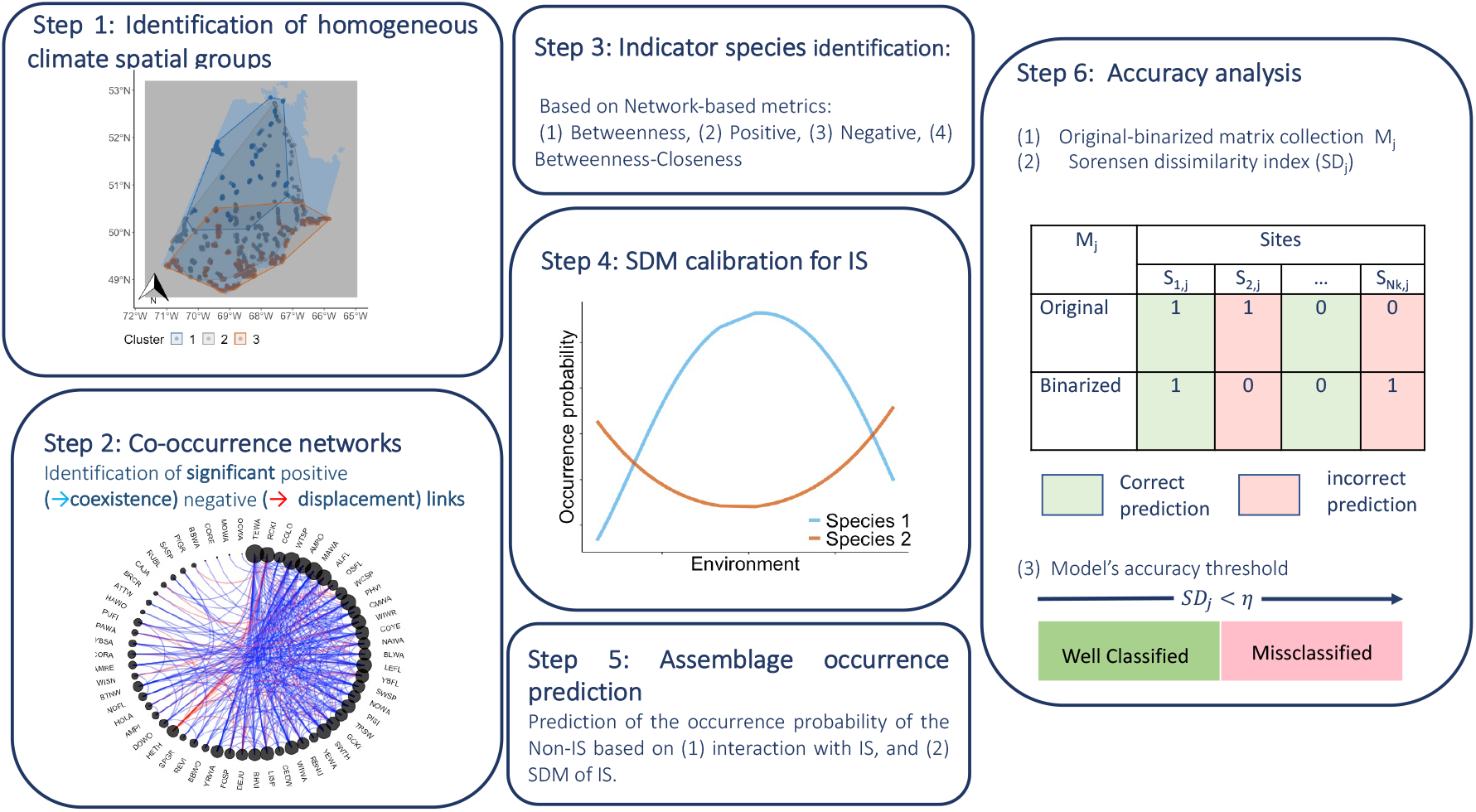
The prediction of species occurrence from IS occurrence involves several steps. In Step 1, we identified spatial clusters based on climate variables. In Step 2, we built co-occurrence networks using Veech’s probabilistic model (Veech, 2013). In Step 3, we identified the IS for each climate region using network-based metrics. In Step 4, we calibrated the Species Distribution Models (SDMs) of the IS for each climate group. In Step 5, we estimated the occurrence probability of non-IS in the pool based on their conditional occurrence relative to the corresponding IS and the occurrence of the IS (Eq. 5). Finally, in Step 6, we assessed the accuracy of our approach through a dissimilarity analysis between the observed occurrences and the estimated ones.

### 2.2 Identification of climate spatial clusters

Static IS are most likely ineffective in monitoring biodiversity over large climate extent (Morelli, 2015; Terrigeol et al., 2022). Therefore, we specified spatial clusters according to a climatic gradient to consider the impact of climate variation on identifying IS and assemblages. We identified the different climate clusters using hierarchical clustering on principal components (package ‘FactoMineR’, Husson et al. (2013)) based on the two first principal components. We used the climate variables as input variables for the principal component analysis (PCA).

### 2.3 Co-occurrence networks

We built a co-occurrence network under each climate cluster to extract the IS. We used occurrence community data (species by site matrix) where 1 indicated a presence and 0 an absence. Species were considered nodes, whereas links reflected positive (coexistence) or negative (displacement) liaisons. To evaluate the cooccurrence significance, we used probabilistic species co-occurrence analysis (Veech, 2013) (‘coccur’ package, Griffith et al. (2016)). Traditionally, building spatial networks of co-occurrence involved principally using null models based on data randomization (Araújo et al., 2011) that is considered a potential source of Type I and Type II errors (Veech, 2013). The probabilistic approach given in Veech (2013) is *distribution-free* and demonstrated to perform better than methods based on data randomization by having a lower Type I error rate and a low Type II error rate. The probability that two species co-occur at *k* sites with max{0, *N*_1_+ *N*_2_− *N*} ≤ *k* ≤ min{N_1_, *N*_2_} followed a hypergeometric distribution 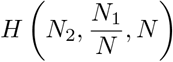 (Veech, 2013):

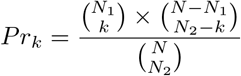

where *N* is the total of sites, *N*_1_ and *N*_2_ are the number of sites occupied by species one and two, respectively.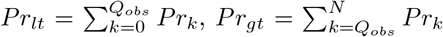 are used as *p*_*values* for testing whether species one and species two co-occur significantly less often or significantly more often, respectively, than expected by chance. *Q*_*obs*_ is the observed number of sites having both species. We included only species pairs with significant positive or negative co-occurrence at *α* = 0.05.

### 2.4 Network-based IS identification

To simplify notations, we remove hereafter the variations between clusters and the method used to extract the IS; however, the procedure remains the same. Our methodology represented each climate cluster by its own set of IS. We considered the number (*q*) of the potential IS (PIS) as well as the number of IS (1 *≤ p < q*). We used four network-based metrics to select the IS belonging to each climate cluster. The first method was based on selecting the first *p* species with maximum significant positive co-occurrences (**Positive**). The second method chose the first *p* species with maximum significant negative co-occurrences (**Negative**). The third method was based on choosing the first *p* species with a maximum normalized betweenness centrality (**Betweenness**):

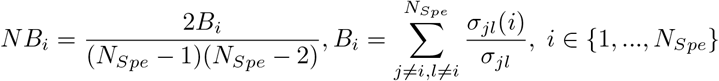

where *σ*_*jl*_(*i*) is the number of the shortest paths relating species j to species l passing by species i and *σ*_*jl*_ is the number of the shortest paths joining species j to species l (we used package ‘igraph’, Csardi and Nepusz (2006) to compute this metric). In the last method, *PIS* (*Spe*_*B*,*q*_) were considered as the *q* species having the maximum of betweenness and the IS will be the most *p* heterogeneous species (maximum normalized closeness centrality) among *Spe*_*B*,*q*_. The betweenness-closeness (**BC**) centrality is given by:

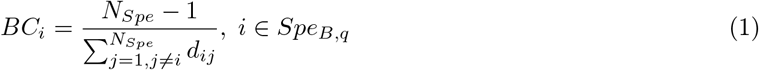

where *d*_*ij*_ is the length of the shortest path between species *i* and *j*. The selection of the IS according to the BC method consists of choosing the first *p* species with minimum BC (Eq. 1). This procedure prevented selecting highly associated species, which most likely represent the same assemblages. (See Fig. 2 for an illustration between IS selected from the four methods).

**Figure 2.**
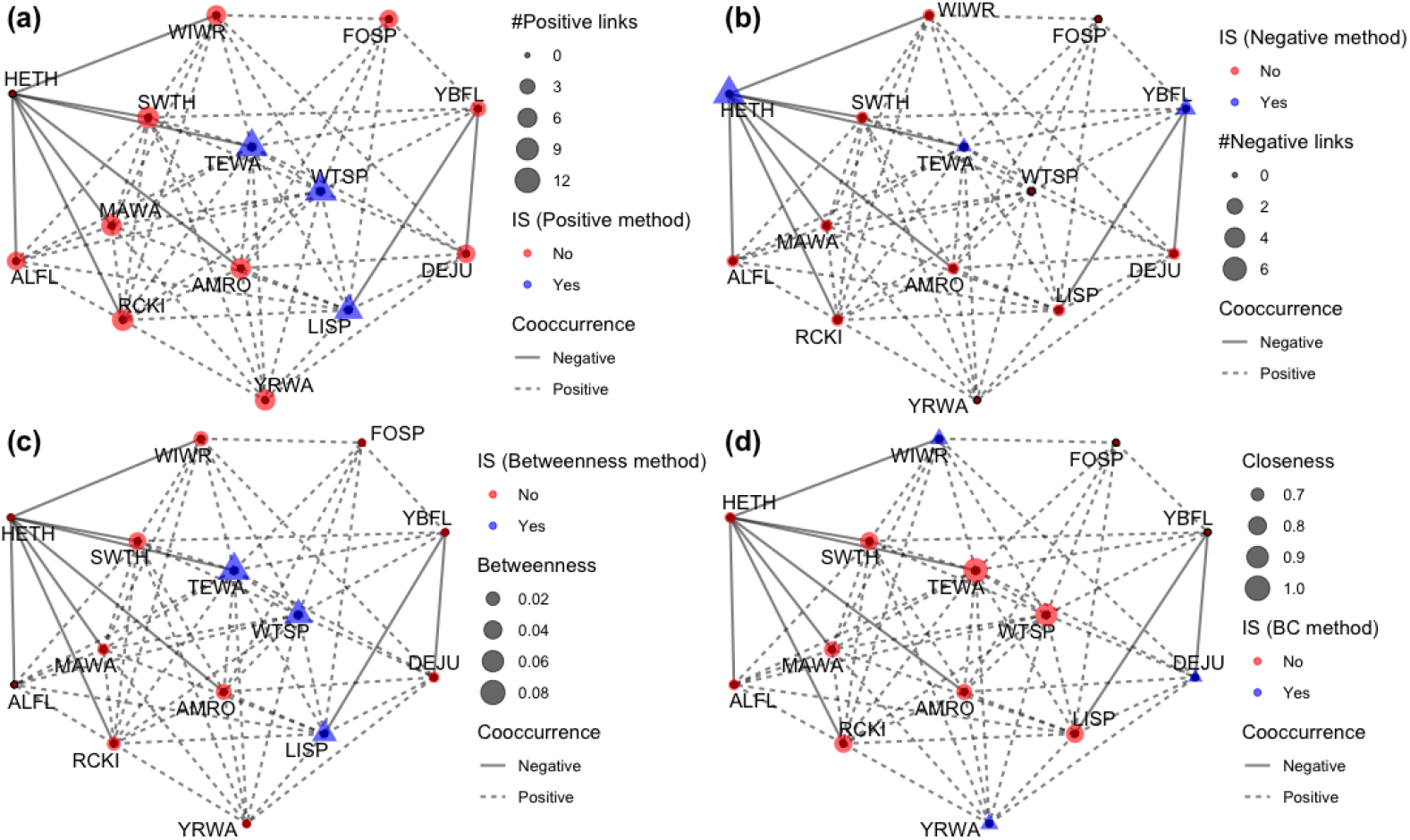
Example of the IS selected from the four metrics. The illustrated co-occurrence network corresponds to the filtered data (at least 20% of presence) for the northern Cluster. The red and blue links represent the negative and the positive significant co-occurrences. The four graphs differ only on the metric for selecting the three IS (triangles). (a) shows the method based on the number of positive c-occurrences, (b) indicates the method based on the number of negative c-occurrences, (c) expresses the method based on the maximum of the betweenness, (d) shows the method based on the minimum of closeness among the PIS having the maximum betweenness (**BC**).

### 2.5 Species distribution models calibration for IS

We calibrated the species distribution models (SDM) of the *jth* IS in two steps: (1) Calibrating the SDM based only on climate variables (*Y*, by including all the potential quadratic and logarithmic transformations) and selecting variables through a step-wise procedure:

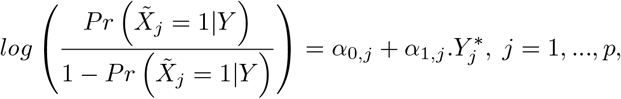

(2) Calibrating the final regression by adding the occurrence of the previous IS as independent variables:

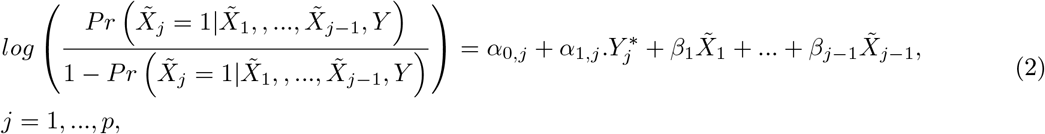

where 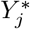 and 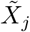 represented respectively the selected predictors and the occurrence of the *jth* IS.

### 2.6 Community occurrence prediction

In our methodology, the occurrence probability of the *jth* non-IS depends implicitly on its interactions with the selected IS set. This probability is a function of (1) the probability of occurrence of IS (Eq. 2) and (2) the probability of presence of the species conditional on the presence or the absence of IS (Eq. 4), which reflects the interaction between the IS and the species in question. The occurrence probability of the *jth* non-IS is given by:

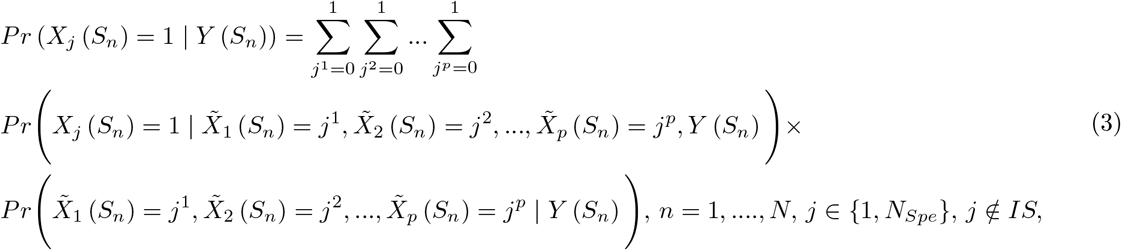

where *X*_*j*_ represents the occurrence of the *jth* non-IS, *S*_*n*_ is the *nth* sampling site and *N* is the total number of sites.

We assume the probability of presence of non-IS conditional to the presence or the absence of the IS and to the environmental condition is invariable spatially:

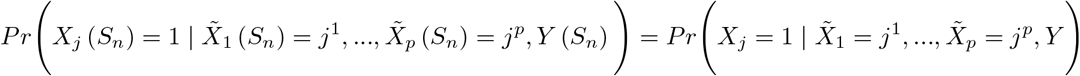

The first component in the right hand of the Eq. 3 is estimated as follows:

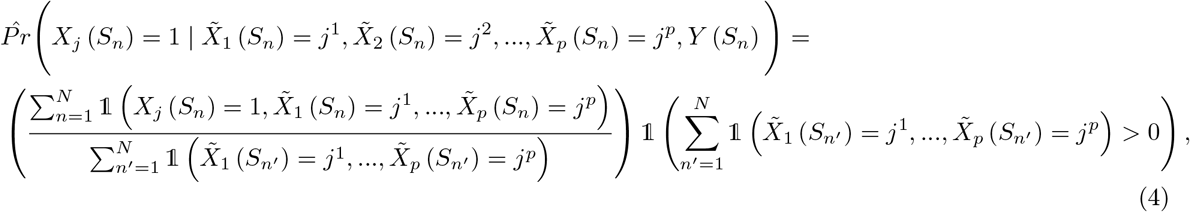

Eq. 2 and Eq. 4 are then used to estimate the occurrence probability of all the other species in the pool:

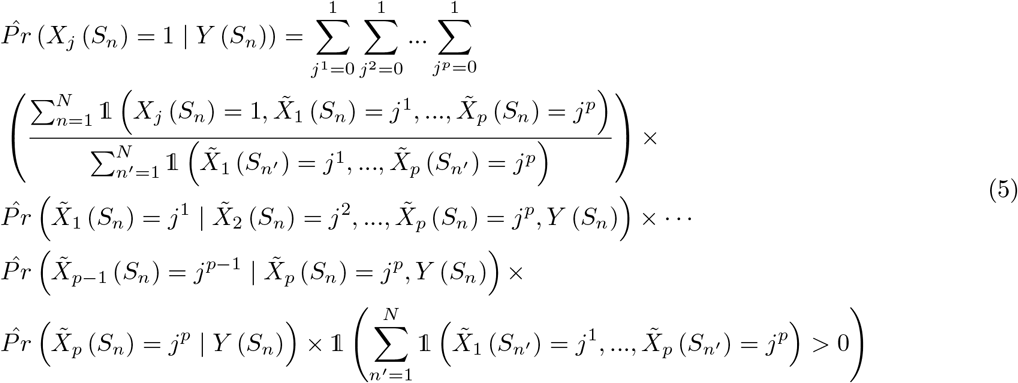

## 3 Accuracy analysis

To determine the binarization threshold for species *j* (*γ*_*j*_), we tested 12 optimization methods aimed at maximizing the accuracy between the actual presence-absence data and the predictions. This included methods such as maximizing the true skill statistic and Kappa statistics (Araújo et al., 2005; Allouche et al., 2006), as implemented in the ‘PresenceAbsence’ package (Freeman and Moisen, 2008). Subsequently, to evaluate the performance of our approach and estimate the number of well-recovered species based on their interaction with IS, we quantified the Sorensen dissimilarity (SD) between the original and the predicted binarized occurrence data:

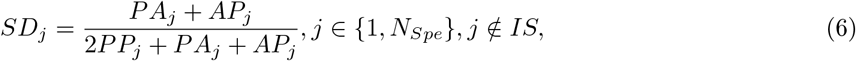

*PP*_*j*_ represents the number of sites where species j is present in both vectors. *PA*_*j*_ represents the number of sites where species j is present in the predicted vector and absent in the observed vector, *AP*_*j*_ the converse. We then studied how the dissimilarity threshold (*η*) influences the percentage of well-classified species (PWS) as a function of the percentage of presence within the dataset. A species is considered ‘well-classified’ if its dissimilarity (*SD*_*j*_) is less than *η*; otherwise, it is considered misclassified. To calculate the percentage of well-recovered species (PWR), we divided the number of well-recovered species by the number of species with significant links within each climate cluster and then multiplied the result by 100.

## 4 Model’s application

### 4.1 Study area and occurrence data

The study area is located in the Côte-Nord region of Québec, Canada (48^°^*N* to 53^°^*N*, 65^°^*W* to 71^°^*W*), within an area of 114118 km2 (Fig. 1). The northern part of the study area belongs to the black spruce-feather moss bioclimatic domain. The southern part of the study area belongs to the eastern balsam fir–white birch subdomain, mostly dominated by balsam fir and white spruce mixed with white (paper) birch.

Presence-absence data were collected between 2010 and 2018 to model species distributions. We used the data from the Atlas of Breeding Birds (Atlas des oiseaux nicheurs du Québec, 2018), which were based on species occurrences that were detected using unlimited distance 5-minute point counts (Bibby et al., 2000), which were collected during the breeding season (late May to mid-July) between 2010 and 2018. To quantify the importance of species occurrence on the performance, we compared the outcomes based on the original presence-absence matrix (**Full data**) and a subset including only species present in more than 20% of sites under each Cluster (**Filtered data**).

### 4.2 Climate predictor variables

To predict species occurrence, we used climate variables. Initially, we generated 17 potential climate variables at a 250-m resolution using the BioSim platform (Régnière et al., 2017), including the annual average of temperature and precipitation between 2004 and 2018 (see Table. 1 in Supporting Information for the description of all potential predictor variables). BioSIM simulates daily maximum and minimum temperatures, precipitation, water deficit, mean daily relative humidity and wind speed by matching georeferenced sources of daily weather data to spatially georeferenced points. BioSIM uses spatial regression to adjust weather data for differences in latitude, longitude, and elevation between the sources of weather data and each field location (for more details, see (Boulanger et al., 2018)). In our case, the spatially referenced points were 15,000 randomly located across the Province of Québec. In contrast, weather data were daily data originating from discrete weather stations within the province. We generated climate variables at a 250 m scale by spatially interpolating data from the 15,000 random points using kriging and elevation as drifting variables.

### 4.3 Identification of climate spatial clusters

We used four climate variables—precipitation, mean temperature, maximum temperature, and minimum temperature—to identify the principal components along the Côte-Nord region in Québec. Hierarchical clustering revealed three climate clusters (see Fig. 3a), indicating a strong latitudinal climatic gradient (see Fig. 3b). This gradient was mainly explained by the latitudinal shift in temperature (see Fig. 3d). For example, Cluster 1, the northernmost cluster, showed the lowest temperature variation with mean temperatures ranging from -4.68°C to -0.28°C. Cluster 2 had mean temperatures between -1.46°C and 1.53°C, while Cluster 3, the southernmost cluster, exhibited the highest temperatures with mean values ranging from 1.31°C to 5.44°C. Precipitation contributed less to the latitudinal explanation of the three climate clusters compared to temperature (see, for instance, the difference between Fig. 3d and Fig. 3e). In this study, we compared only Clusters 1 and 3 (see S2 File for the data under the two climate clusters).

**Figure 3.**
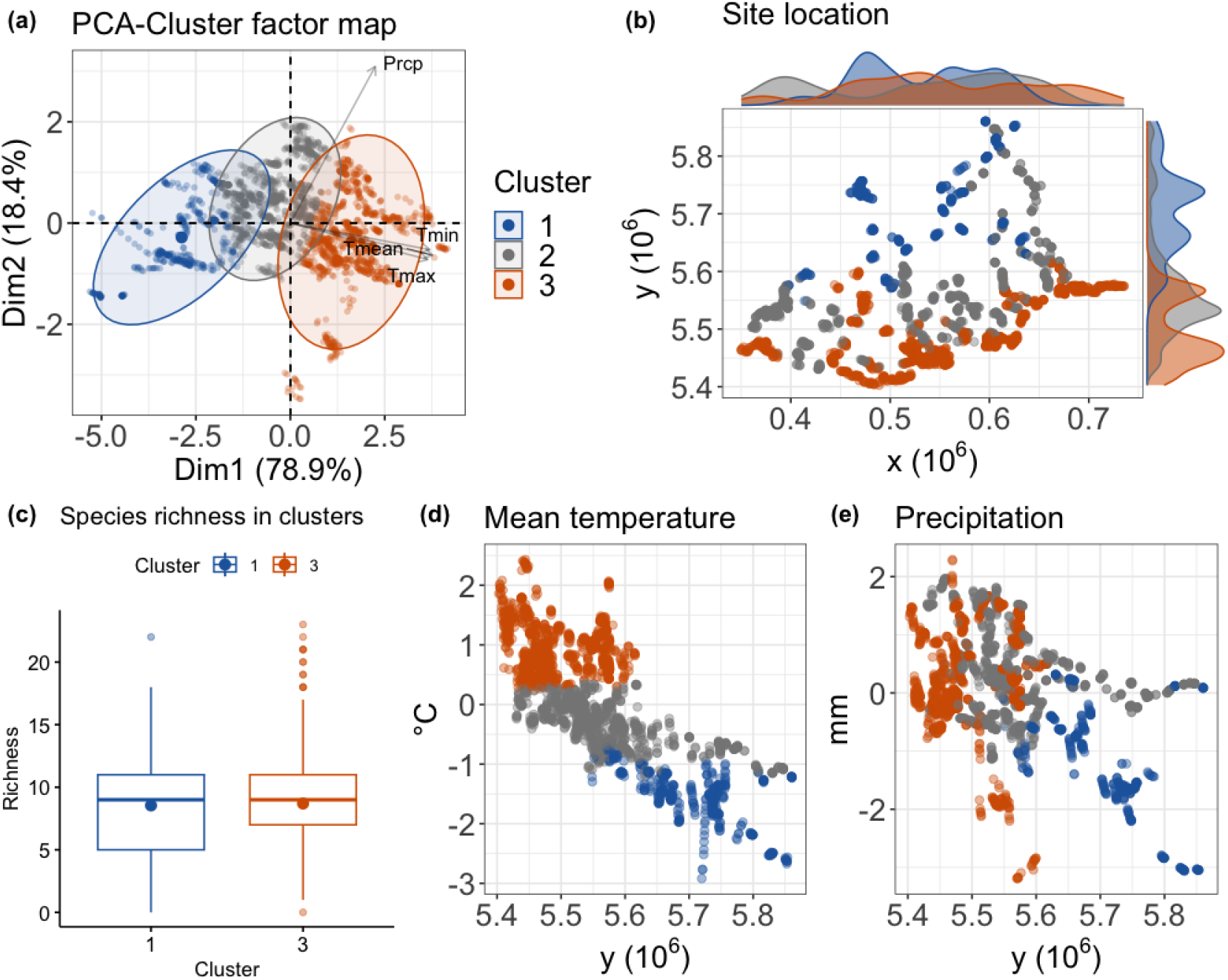
Identification of climate spatial clusters. (a) shows the factor map of Principal Component Analysis (PCA). The hierarchical clustering is applied to the PCA, and the results allowed three distinct homogeneous latitudinal climate clusters: Cluster 1 is northern, and Cluster 3 is southern. (b) The spatial distribution of the three climate clusters. (c) Species richness at the northern and southern clusters. (d)-(e) The distribution of temperature and precipitation within the three climate clusters.

### 4.4 Co-occurrence network analysis

We considered each climate cluster as a network and analyzed the outputs of the northern (Cluster 1) and southern (Cluster 3) co-occurrence networks. Cluster 3 included more species than Cluster 1 (88 versus 61 species, respectively), with more than double the number of sites (979 versus 406, respectively). However, Cluster 3, the southern cluster, exhibited nearly double the percentage of negative co-occurrence links compared to the northern cluster (25.2% versus 13.2%, respectively; see Fig. 4b). Additionally, more species had higher co-occurrences (with a degree between 20 and 30) in the south than in the north, whereas more species had low co-occurrences (with a degree between 0 and 20) in the north than in the south (see Fig. 4c). Furthermore, we observed a bimodal degree distribution in both networks, indicating two groups of weak-interacting and strong-interacting species. The separation between weak-interacting and strong-interacting species was more pronounced in the northern cluster, with a sharper distinction between the two groups under Cluster 1 (see Fig. 4c).

**Figure 4.**
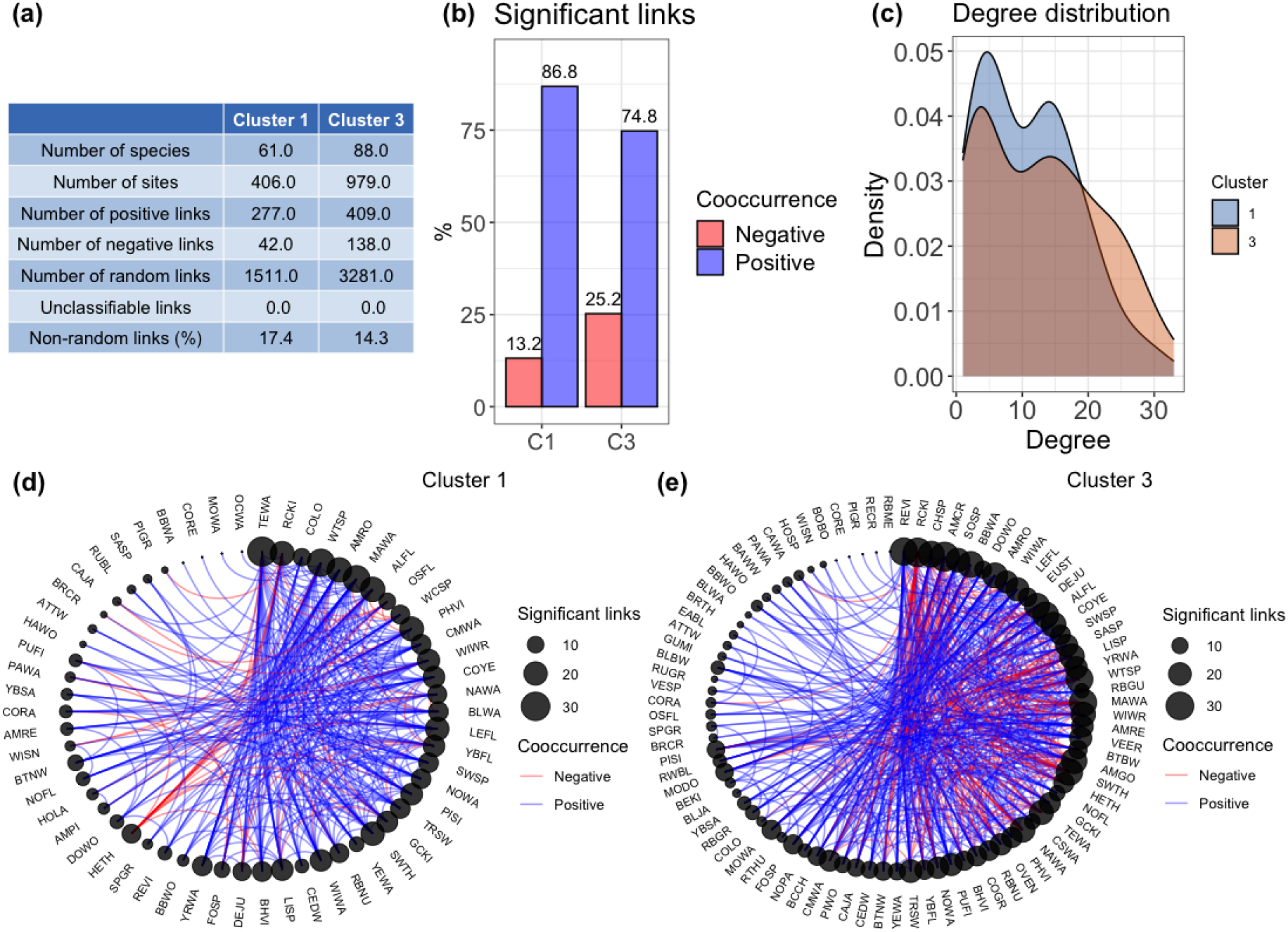
Networks characteristics. (a) The information related to the northern (Cluster 1) and southern (Cluster 3) networks. (b) The degree distribution of each network. (c) The percentage of positive and negative links (co-occurrences) within the northern (C1) and southern (C3) networks. (d)-(e) The northern and southern co-occurrence networks. The blue and red links represent the negative and positive significant co-occurrences. The size of the circle for each species represents the degree.

### 4.5 Network-based indicator species

We identified three influential species (IS) based on four network metrics (“Positive”, “Negative”, “Betweenness”, and “Betweenness-Closeness”) for the northern (Cluster 1) and southern (Cluster 3) climate networks. Additionally, we included a fifth metric representing the “combined” case, selecting the best metric for each species that minimized the dissimilarity between the original and estimated occurrence. For each of the four network-based metrics, we selected the three most informative species.

When comparing latitudinal variation in IS, only two species remained common IS between the northern and southern networks among the selected 17 IS. The other 15 species were either unique to the northern or southern networks (see Fig. 5a). For both networks, at most one IS was shared among the three species selected for each metric (see Fig. 5b-c). A similarity of 1 implies that the three IS are the same and a value of 0 implies that the three IS are different between the two metrics. Notably, IS selected based on betweenness centrality were most similar to those selected using the other metrics. In Cluster 1 (northern network), the similarity between IS based on betweenness centrality and those based on betweenness-closeness centrality and positive co-occurrences was 0.33. In Cluster 3 (southern network), the similarity between IS based on betweenness centrality and those based on positive and negative co-occurrences was also 0.33. Betweenness-closeness centrality was similar to betweenness centrality and negative co-occurrence only in the northern network. Overall, the similarity in IS selection was higher in the northern network than in the southern network. In the northern network, each metric shared at least one IS with another metric, whereas in the southern network, “BC” did not share any common IS with other metrics.

**Figure 5.**
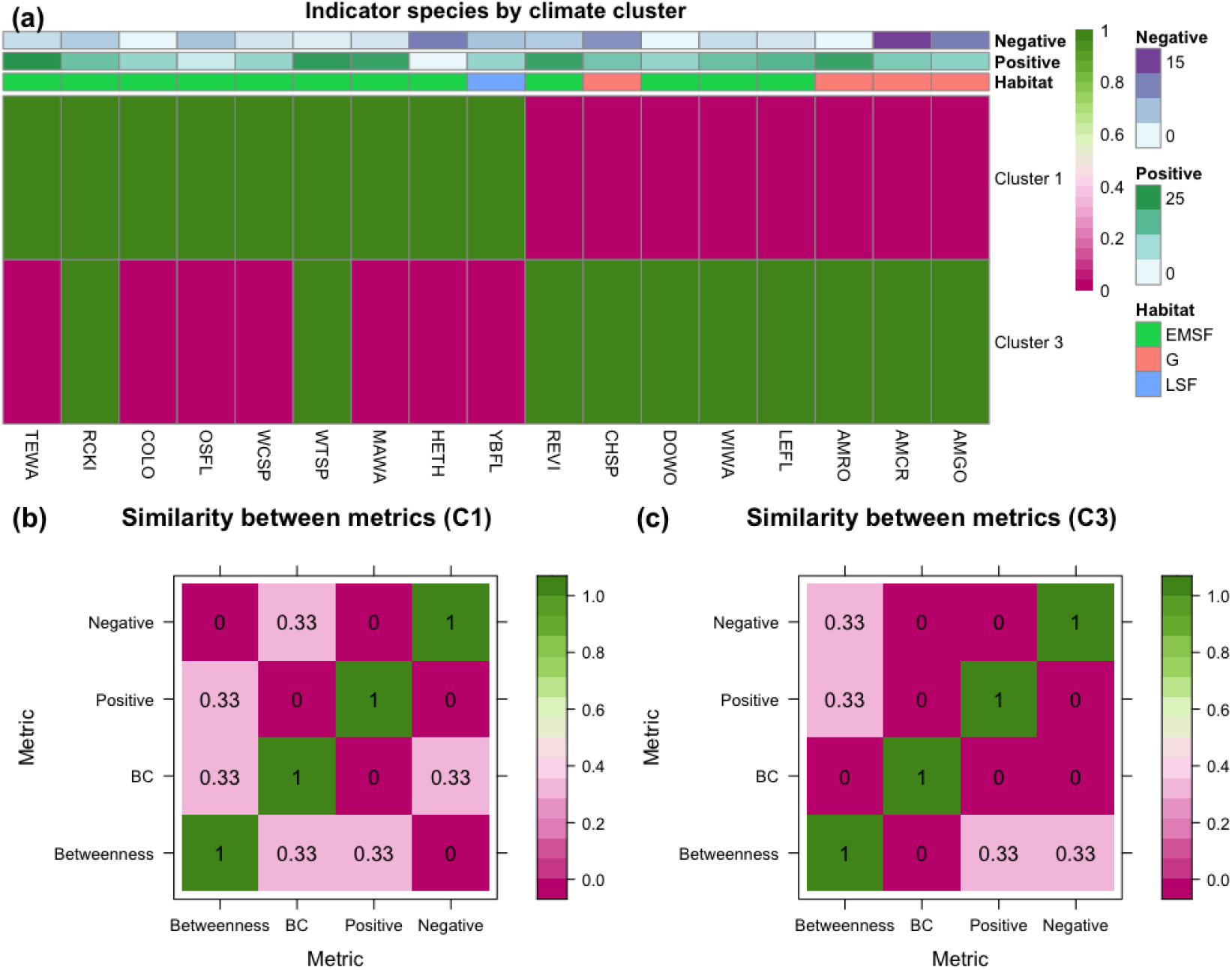
The similarity in the selection of IS is based on the four metrics: betweenness-closeness, betweenness, negative co-occurrences and positive co-occurrences within each of the northern and the southern climate clusters. (a) The identified IS according to all the metrics. Values of one and zero indicate that the species is either an indicator or not an IS for the specified climate cluster. We also showed the species’ habitat and the number of positive and negative significant co-occurrences for each species. Habitat shows the IS habitat: late succession forest (*LSF*), early-to-mid succession forest (*EMSF*) or generalist species (*G*) (see S1 and S1 File). (b)-(c) Similarity with a value of 0 indicates no common IS between a pair of metrics. A value of 1 indicates the same set of IS between the two metrics (see S1 Table).

A significant difference in the habitat of the IS between the northern and southern networks was observed (see S1). Although most IS were associated with early-to-mid succession forests, only one species, the *Yellow-bellied Flycatcher* (YBFL), was associated with late succession forests, and this occurred only in the northern network. All four generalist species were selected as IS exclusively within the southern network (see S1 Table).

### 4.6 Performance of the indicator species in recovering assemblages occurrence

According to all metrics, there is a negative relationship between species presence percentage and the dissimilarity between the true and estimated occurrences (see Fig. 6). Specifically, the Pearson correlation between SD and the percentage of presence for Cluster 1 ranged from -0.97 (based on the betweenness-closeness method) to -0.95 (based on the Positive method). For Cluster 3, the correlation between SD and the percentage of presence ranged from -0.98 (based on the betweenness method) to -0.97 (for all other methods). Therefore, species with higher presence percentages are more efficiently recovered based on their co-occurrence with IS. In addition, by comparing the two networks, we observed that the percentage of presence must be significantly higher in the southern network than in the northern network to achieve an SD less than 0.5. The minimum percentage of presence required for *η* = 0.49 was 29.06% for the northern network, while for *η* = 0.50, the corresponding value was 33.20% for the southern network (see Fig. 6) To evaluate how the percentage of presence impacted the effectiveness in estimating species occurrence, we compared the percentage of well-recovered species between full and filtered data for the combined metrics case (see Fig. 7). A species was considered well-recovered if the Sorensen dissimilarity (*SD*) between the original and estimated occurrences was less than *η* = 0.5. The filtered data included only species with a presence percentage of at least 20%. For the filtered data, the percentage of well-recovered species was 85.71% within the northern network (Cluster 1) and 53.84% within the southern network (Cluster 3). In contrast, for the full data, the percentage of well-recovered species was 27.59% within the northern network (Cluster 1) and 9.41% within the southern network (Cluster 3).

**Figure 6.**
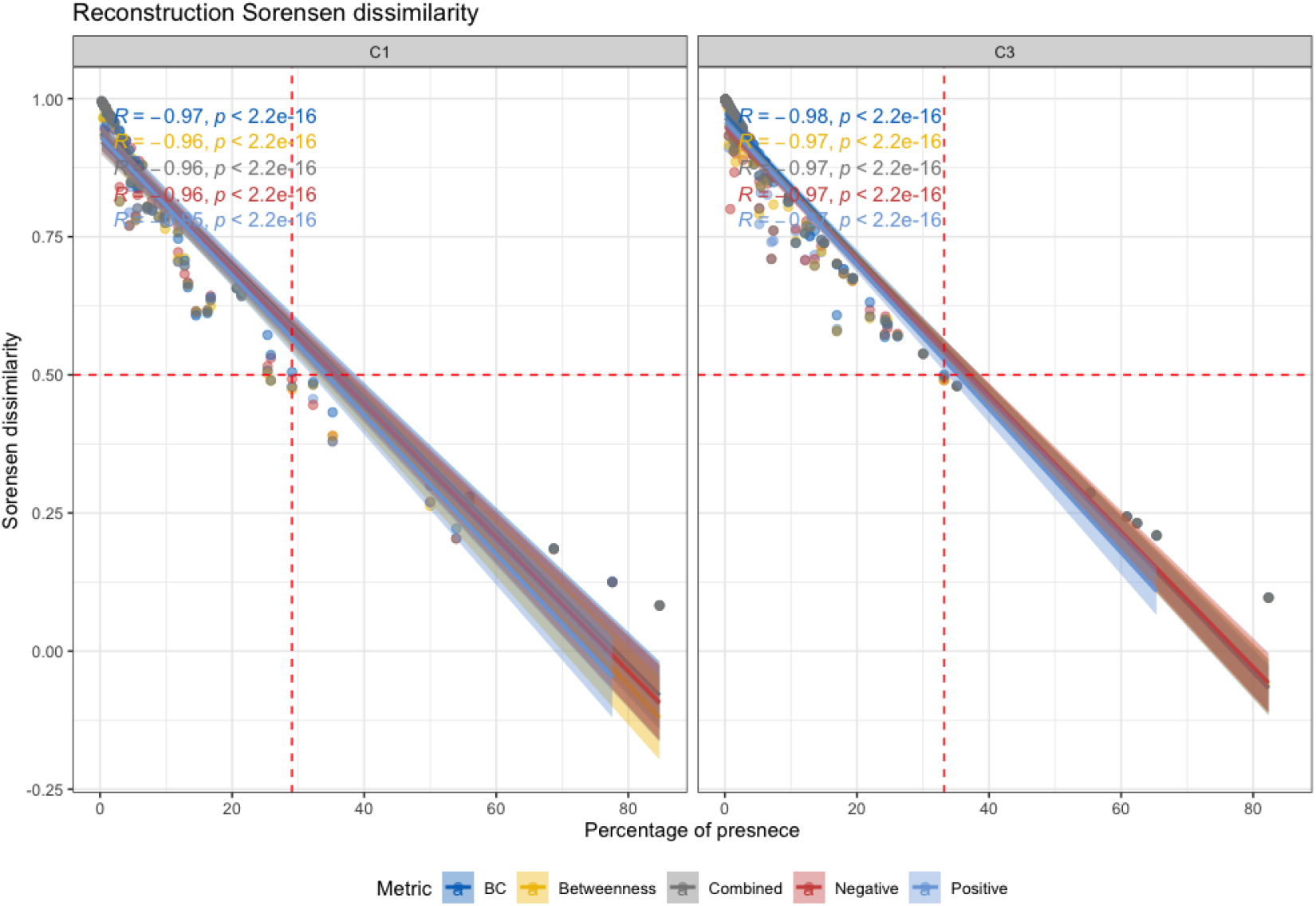
Cluster assemblage Sorensen dissimilarity (SD) vs. species’ percentage of presence within the northern (C1) and southern (C3) climate clusters.

**Figure 7.**
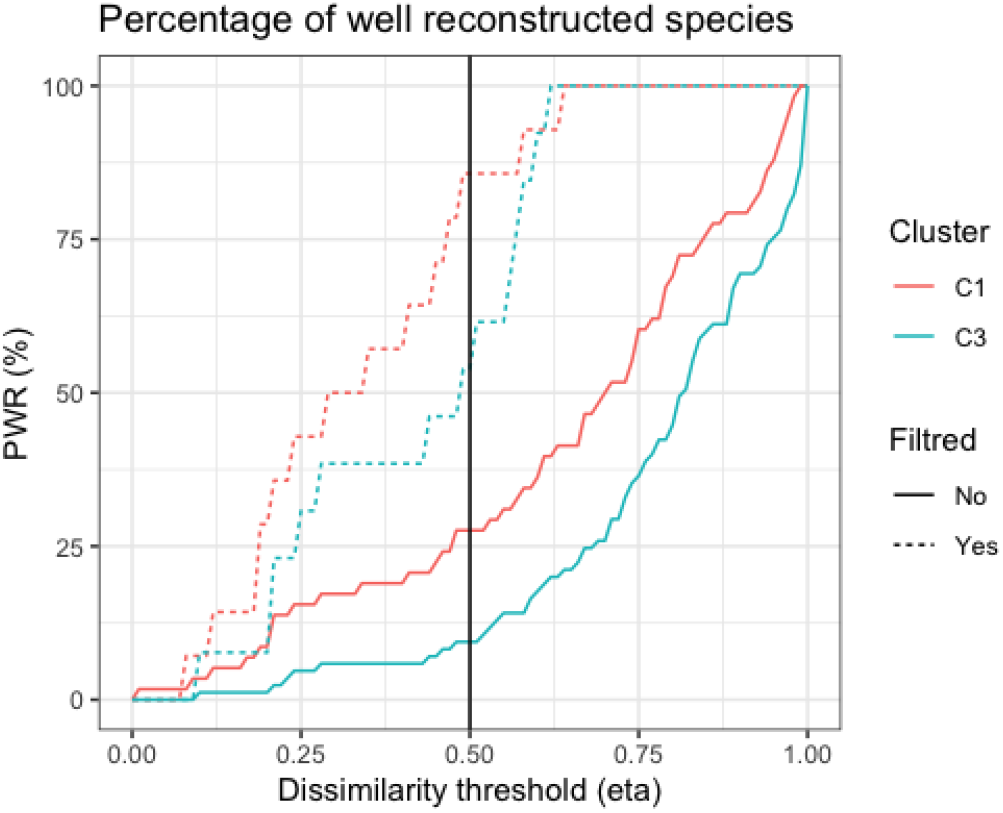
Percentage of well-recovered species using the ‘combined’ method, based on the dissimilarity between original and predicted occurrences. A species is considered well-recovered if *η* = 0.5. Analysis was performed on both the full dataset (non-filtered) and a filtered dataset, were species present in less than 20% of sites were excluded.

## 5 Discussion

We proposed a network-based methodology to assess the quality of biodiversity reconstruction based on co-occurrences with IS, which serves as a informative tool for monitoring biodiversity (Elmendorf et al., 2015; Mimet et al., 2016). This study aimed to quantify the informativeness of IS in reconstructing species assemblages and to determine the conditions under which this reconstruction is most effective. We were based on network-based IS identification followed by co-occurrence network analysis and species distribution modeling in order to calculate the probability of occurrence of the assemblage along a latitudinal network. We implicitly incorporated the significant co-occurrences between IS and the assemblage by computing the probability of species occurrence conditional on the presence or the absence of IS. One key finding is that the more a species is present, the better the quality of biodiversity reconstruction. Also, the biodiversity reconstruction is negatively related to the inter-specific competition which higher toward the south. However, the conditions for effective recovery of assemblage occurrence become more severe towards lower latitudes. This implies that species must be more present under latitudes closer to Tropics to be well recovered from their co-occurrences with IS.

### Latitudinal gradient and the alternation in IS composition and interspecific interactions

Latitudinal data offers the almost unique possibility of studying climate change’s effect on the composition of assemblages. The latitudinal networks consist of arrays along the latitude axis representing co-occurring species (Pérez-García et al., 2016) within a homogeneous climate extent (Terrigeol et al., 2022). Anthropogenic climate change can alter interspecific interactions and produce unexpected changes in species distributions, community structure, and diversity (Harley, 2011). Accordingly, we noticed that the latitudinal gradient plays a crucial role in the change in the nature of the composition of the IS and the biotic interactions. Indeed, there is an almost complete turnover in the composition of the IS between the northern and southern networks based on the network-based metrics. This observed change in the composition of the IS is in line with other methods based on species richness (Terrigeol et al., 2022). Consequently, short-term climate change could lead to different IS for species assemblage. Also, Blanchet et al. (2020) showed that often, species co-occur without significant interaction, and consequently, it becomes more complicated to belong to a shared community under different climate conditions.

Furthermore, by comparing the northern and the southern networks, we predict a rise in the percentage of the negative co-occurrence liaisons toward the south. Indeed, the percentage of negative co-occurrences increased from 13.2% to 25.2% from the northern to the southern network. Recent observations showed that climate change can change the abundance of species and alter biotic interaction with a loss of interactions through extinctions, range shifts, and changes in relative abundance (Blois et al., 2013). Accordingly, Milazzo et al. (2013) showed that species co-occurrence could switch to competitive displacement mode in rising temperatures. A clear shift in the temperature was noticed with a difference that reaches more than 6*°*C between the northern and the southern networks (see Fig. 3), which could be a fundamental reason for this change (Milazzo et al., 2013). Accordingly, the rise in the temperature could increase the competition and displacement could also lead to the switch to a competitive displacement mode (Milazzo et al., 2013) or to an increase in the predation intensity (Harley, 2011). A study by DeGregorio et al. (2015) demonstrated that climate change is expected to increase predation of bird nests by ratsnakes by 7%. Moreover, Delavaux et al. (2024) showed evidence that mutualisms, which is a positive interaction, is one of keys to the maintenance of high tropical plant diversity and mediate the biogeographic patterns of plant diversity on Earth. Specifically, only one identified IS belonging to late succession forest habitat all the 16 other species were either generalist or belonging to early-to mid succession forest habitat.

### The effectiveness in the reconstruction of assemblage occurrence

In order to reconstruct biodiversity based on the co-occurrences with IS, the percentage of presence must be at least 29% for the northern community and 33% for the southern community to reach a dissimilarity less than 0.5. Therefore, it becomes challenging to recover the rare species. This could be due to the spatial variation in species assemblages. The complex interactions within ecosystems can make it difficult to reliably infer the status of rare species from the presence or absence of IS, particularly when those indicators are generalist species that may thrive under a wide range of conditions (Hilty and Merenlender, 2000).

In addition, it is more challenging to reconstruct biodiversity in the southern community due to increased competition between species impacting the co-occurrences (Milazzo et al., 2013). On one hand, the higher levels of biodiversity in tropical regions may pose challenges for reconstruction, as the complex interactions and interspecific competition in these areas can make it more difficult to infer community composition from species co-occurrences. On the other hand, the lower minimum requirement of the percentage of presence at the north sites reflects the lower species richness observed at higher latitudes (see Fig. 3c). This pattern could be explained by the latitudinal diversity gradient where the richness increases toward the Tropics (Pianka, 1966; Rohde, 1992; Hillebrand, 2004).

## Conclusion

We used network-based framework to study the patterns and extent of changes in interactions between species latitudinaly. The framework provided an estimation of the likelihood of species’ occurrences based on their implicit interactions with local IS, which is most effective for species that are highly abundant. This approach quantifies the potential changes in species’ interactions due to climate change. However, the limited success, particularly in recovering rare species, highlights the need for further theoretical investigation of the modeling framework. Interestingly, the latitudinal gradient significantly influences biotic interactions and the composition of species groups, incorporating a latitudinal network could help describing interactions between different layers, such as dispersal and changes in the composition of species groups along the latitudinal axis. Using multiplex complex network analysis to explore interactions between networks, as suggested by (Lurgi et al., 2020), could enhance the model.

## Acknowledgements

This work was supported by the Sentinel North programme of Laval University, funded by the Canada First Research Excellence Fund. AA, DF and PD were also supported by the Natural Sciences and Engineering Research Council of Canada (NSERC). We acknowledge Calcul Québec and Compute Canada for their technical support and computing infrastructures. We thank also the Québec Breeding Bird Atlas for supplying data. We would also like to thank the following partners: Regroupement QuébecOiseaux, Environment and Climate Change Canada and Birds Canada, together with all the volunteer participants who gathered data for the project. We are grateful to Louis-Paul Rivest for his valuable statistical suggestions and advice.

## Supporting information

**S1 Habitat types of bird species.**

**S1 Table Network-based indicator species.**

**S1 File The associated habitat of the potential bird species.**

**S2 File The occurrence data for the northern and southern climate clusters.**

## References

Allouche, O., Tsoar, A., and Kadmon, R. (2006). Assessing the accuracy of species distribution models: prevalence, kappa and the true skill statistic (TSS). Journal of Applied Ecology, 43(6):1223–1232.

Araújo, M. B., Rozenfeld, A., Rahbek, C., and Marquet, P. A. (2011). Using species co-occurrence networks to assess the impacts of climate change. Ecography, 34(6):897–908.

Araújo, M. B., Pearson, R. G., Thuiller, W., and Erhard, M. (2005). Validation of species–climate impact models under climate change. Global Change Biology, 11(9):1504–1513.

Atlas des oiseaux nicheurs du Québec (2018). Données obtenues en réponse à une demande présentée aux bureaux de l’Atlas (www.atlas-oiseaux.qc.ca). Regroupement QuébecOiseaux, Service canadien de la faune d’Environnement Canada et Études d’Oiseaux Canada. Québec, QC, Canada.

Azeria, E. T., Fortin, D., Hébert, C., Peres-Neto, P., Pothier, D., and Ruel, J.-C. (2009). Using null model analysis of species co-occurrences to deconstruct biodiversity patterns and select indicator species. Diversity and Distributions, 15(6):958–971.

Bibby, C. J., Burgess, N. D., Hill, D. A., Hillis, D. M., and Mustoe, S. (2000). Bird census techniques. Elsevier.

Blanchet, F. G., Cazelles, K., and Gravel, D. (2020). Co-occurrence is not evidence of ecological interactions. Ecology Letters, 23(7):1050–1063.

Blois, J. L., Zarnetske, P. L., Fitzpatrick, M. C., and Finnegan, S. (2013). Climate change and the past, present, and future of biotic interactions. Science, 341(6145):499–504.

Bouderbala, I., Labadie, G., Béland, J.-M., Boulanger, Y., Hébert, C., Desrosiers, P., Allard, A., and Fortin, D. (2023a). Effects of global change on bird and beetle populations in boreal forest landscape: An assemblage dissimilarity analysis. Diversity and Distributions, 29(6):757–773.

Bouderbala, I., Labadie, G., Béland, J.-M., Tremblay, J. A., Boulanger, Y., Hébert, C., Desrosiers, P., Allard, A., and Fortin, D. (2023b). Long-term effect of forest harvesting on boreal species assemblages under climate change. PLOS Climate, 2(3):1–20.

Boulanger, Y., Parisien, M.-A., and Wang, X. (2018). Model-specification uncertainty in future area burned by wildfires in Canada. International Journal of Wildland Fire, 27(3):164–175.

Boulanger, Y. and Puigdevall, J. P. (2021). Boreal forests will be more severely affected by projected anthropogenic climate forcing than mixedwood and northern hardwood forests in eastern Canada. Landscape Ecology, 36(6):1725–1740.

Boulanger, Y., Taylor, A. R., Price, D. T., Cyr, D., McGarrigle, E., Rammer, W., Sainte-Marie, G., Beaudoin, A., Guindon, L., and Mansuy, N. (2017). Climate change impacts on forest landscapes along the Canadian southern boreal forest transition zone. Landscape Ecology, 32(7):1415–1431.

Cadieux, P., Boulanger, Y., Cyr, D., Taylor, A. R., Price, D. T., Sólymos, P., Stralberg, D., Chen, H. Y., Brecka, A., and Tremblay, J. A. (2020). Projected effects of climate change on boreal bird community accentuated by anthropogenic disturbances in western boreal forest, Canada. Diversity and Distributions, 26(6):668–682.

Chu, T.-J., Shih, C.-H., Lu, Y.-M., Shih, Y.-J., Wang, J.-Q., and Huang, L.-M. (2021). Incorporating species-conditional co-occurrence when selecting indicator species to monitor restoration after mangrove removal from the siangshan wetland, taiwan. Journal of Marine Science and Engineering, 9(10).

Csardi, G. and Nepusz, T. (2006). The igraph software package for complex network research. InterJournal, Complex Systems:1695.

DeGregorio, B. A., Westervelt, J. D., Weatherhead, P. J., and Sperry, J. H. (2015). Indirect effect of climate change: Shifts in ratsnake behavior alter intensity and timing of avian nest predation. Ecological Modelling, 312:239–246.

Delavaux, C. S., Crowther, T. W., Bever, J. D., Weigelt, P., and Gora, E. M. (2024). Mutualisms weaken the latitudinal diversity gradient among oceanic islands. Nature, pages 1–13.

Dommain, R., Andama, M., McDonough, M. M., Prado, N. A., Goldhammer, T., Potts, R., Maldonado, J. E., Nkurunungi, J. B., and Campana, M. G. (2020). The challenges of reconstructing tropical biodiversity with sedimentary ancient dna: A 2200-year-long metagenomic record from bwindi impenetrable forest, uganda. Frontiers in Ecology and Evolution, 8.

Dufrêne, M. and Legendre, P. (1997). Species assemblages and indicator species:the need for a flexible asymmetrical approach. Ecological Monographs, 67(3):345–366.

Duveneck, M. J., Scheller, R. M., White, M. A., Handler, S. D., and Ravenscroft, C. (2014). Climate change effects on northern Great Lake (USA) forests: A case for preserving diversity. Ecosphere, 5(2):art23.

Elmendorf, S. C., Henry, G. H. R., Hollister, R. D., Fosaa, A. M., Gould, W. A., Hermanutz, L., Hofgaard, A., Jónsdóttir, I. S., Jorgenson, J. C., Lévesque, E., Magnusson, B., Molau, U., Myers-Smith, I. H., Oberbauer, S. F., Rixen, C., Tweedie, C. E., and Walker, M. D. (2015). Experiment, monitoring, and gradient methods used to infer climate change effects on plant communities yield consistent patterns. Proceedings of the National Academy of Sciences, 112(2):448–452.

Fleming, J., Sutherland, C., Sterrett, S. C., and Campbell Grant, E. H. (2020). A latent process model approach to improve the utility of indicator species. Oikos, 129(12):1753–1762.

Freeman, E. A. and Moisen, G. (2008). PresenceAbsence: An r package for presence absence analysis. Journal of Statistical Software, 23(11):1–31.

Gotelli, N. J. (2000). Null model analysis of species co-occurrence patterns. Ecology, 81(9):2606–2621.

Graham, R. W. and Grimm, E. C. (1990). Effects of global climate change on the patterns of terrestrial biological communities. Trends in Ecology and Evolution, 5(9):289–292.

Griffith, D. M., Veech, J. A., and Marsh, C. J. (2016). Cooccur: probabilistic species co-occurrence analysis in r. Journal of Statistical Software, 69:1–17.

Harley, C. D. G. (2011). Climate change, keystone predation, and biodiversity loss. Science, 334(6059):1124– 1127.

Hillebrand, H. (2004). On the generality of the latitudinal diversity gradient. The American Naturalist, 163(2):192–211. PMID: 14970922.

Hillebrand, H., Soininen, J., and Snoijs, P. (2010). Warming leads to higher species turnover in a coastal ecosystem. Global Change Biology, 16(4):1181–1193.

Hilty, J. and Merenlender, A. (2000). Faunal indicator taxa selection for monitoring ecosystem health. Biological Conservation, 92(2):185–197.

Husson, F., Josse, J., Le, S., and Mazet, J. (2013). Factominer: multivariate exploratory data analysis and data mining with r. R package version, 1(1.29).

Kelly, A. E. and Goulden, M. L. (2008). Rapid shifts in plant distribution with recent climate change. Proceedings of the National Academy of Sciences of the United States of America, 105(33):11823–11826.

Kim, K. C. and Byrne, L. B. (2006). Biodiversity loss and the taxonomic bottleneck: emerging biodiversity science. Ecological Research, 21(6):794.

Labadie, G., Hardy, C., Boulanger, Y., Vanlandeghem, V., Hebblewhite, M., and Fortin, D. (2023). Global change risks a threatened species due to alteration of predator–prey dynamics. Ecosphere, 14(3):e4485.

Labadie, G., McLoughlin, P. D., Hebblewhite, M., and Fortin, D. (2021). Insect-mediated apparent competition between mammals in a boreal food web. Proceedings of the National Academy of Sciences of the United States of America, 118(30).

Lurgi, M., Galiana, N., Broitman, B. R., Kéfi, S., Wieters, E. A., and Navarrete, S. A. (2020). Geographical variation of multiplex ecological networks in marine intertidal communities. Ecology, 101(11):e03165.

Masson-Delmotte, V., Zhai, P., Pörtner, H.-O., Roberts, D., Skea, J., Shukla, P. R., et al. (2018). Global Warming of 1.5 ?C. An IPCC Special Report on the impacts of global warming of 1.5 ?C above pre-industrial levels and related global greenhouse gas emission pathways, in the context of strengthening the global response to the threat of climate change, sustainable development, and efforts to eradicate poverty. Technical report, IPCC.

Micheletti, T., Stewart, F. E. C., Cumming, S. G., Haché, S., Stralberg, D., Tremblay, J. A., Barros, C., Eddy, I. M. S., Chubaty, A. M., Leblond, M., Pankratz, R. F., Mahon, C. L., Van Wilgenburg, S. L., Bayne, E. M., Schmiegelow, F., and McIntire, E. J. B. (2021). Assessing pathways of climate change effects in SpaDES: An application to bBoreal Landbirds of Northwest Territories Canada. Frontiers in Ecology and Evolution, 9(October).

Milazzo, M., Mirto, S., Domenici, P., and Gristina, M. (2013). Climate change exacerbates interspecific interactions in sympatric coastal fishes. Journal of Animal Ecology, 82(2):468–477.

Mimet, A., Pellissier, V., Houet, T., Julliard, R., and Simon, L. (2016). A holistic landscape description reveals that landscape configuration changes more over time than composition: Implications for landscape ecology studies. PLOS ONE, 11(3):1–16.

Morelli, F. (2015). Indicator species for avian biodiversity hotspots: Combination of specialists and generalists is necessary in less natural environments. Journal for Nature Conservation, 27:54–62.

Pachauri, R. K., Allen, M. R., Barros, V. R., Broome, J., Cramer, W., Christ, R., Church, J. A., Clarke, L., Dahe, Q., Dasgupta, P., et al. (2014). Climate change 2014: synthesis report. Contribution of Working Groups I, II and III to the fifth assessment report of the Intergovernmental Panel on Climate Change. IPCC, Gland, Switzerland.

Parmesan, C. and Yohe, G. (2003). A globally coherent fingerprint of climate change impacts across natural systems. Nature, 421(6918):37–42.

Pianka, E. R. (1966). Latitudinal gradients in species diversity: A review of concepts. The American Naturalist, 100(910):33–46.

Pérez-García, J. M., Sebastián-González, E., Botella, F., and Sánchez-Zapata, J. A. (2016). Selecting indicator species of infrastructure impacts using network analysis and biological traits: Bird electrocution and power lines. Ecological Indicators, 60:428–433.

Rechkemmer, A. and von Falkenhayn, L. (2009). The human dimensions of global environmental change: Ecosystem services, resilience, and governance.

Régnière, J., Saint-Amant, R., Béchard, A., and Moutaoufik, A. (2017). BioSIM 11: User’s manual. Technical report, Natural Resources Canada, Canadian Forest Service, Laurentian Forestry Centre, Québec, QC Canada.

Rivest, L.-P. and Ebouele, S. E. (2020). Sampling a two dimensional matrix. Computational Statistics Data Analysis, 149:106971.

Robichaud, C. B. and Knopff, K. H. (2015). Biodiversity offsets and caribou conservation in alberta: opportunities and challenges. Rangifer, 35(2):99–122.

Rohde, K. (1992). Latitudinal gradients in species diversity: The search for the primary cause. Oikos, 65(3):514–527.

Santillán, V., Quitián, M., Tinoco, B. A., Zárate, E., Schleuning, M., Böhning-Gaese, K., and Neuschulz, E. L. (2018). Spatio-temporal variation in bird assemblages is associated with fluctuations in temperature and precipitation along a tropical elevational gradient. PLOS ONE, 13(5):1–15.

Siddig, A. A., Ellison, A. M., Ochs, A., Villar-Leeman, C., and Lau, M. K. (2016). How do ecologists select and use indicator species to monitor ecological change? insights from 14 years of publication in ecological indicators. Ecological Indicators, 60:223–230.

Stralberg, D., Bayne, E. M., Cumming, S. G., Sólymos, P., Song, S. J., and Schmiegelow, F. K. (2015). Conservation of future boreal forest bird communities considering lags in vegetation response to climate change: A modified refugia approach. Diversity and Distributions, 21(9):1112–1128.

Terrigeol, A., Ewane Ebouele, S., Darveau, M., Hébert, C., Rivest, L.-P., and Fortin, D. (2022). On the efficiency of indicator species for broad-scale monitoring of bird diversity across climate conditions. Ecological Indicators, 137:108773.

Thuiller, W., Lavergne, S., Roquet, C., Boulangeat, I., Lafourcade, B., and Araujo, M. B. (2011). Consequences of climate change on the tree of life in Europe. Nature, 470(7335):531–534.

Veech, J. A. (2013). A probabilistic model for analysing species co-occurrence. Global Ecology and Biogeog-raphy, 22(2):252–260.

Zhang, Y. and Liang, S. (2014). Changes in forest biomass and linkage to climate and forest disturbances over northeastern china. Global Change Biology, 20(8):2596–2606.

